# Recent progress in multi-electrode spike sorting methods

**DOI:** 10.1101/086991

**Authors:** Baptiste Lefebvre, Pierre Yger, Olivier Marre

## Abstract

In recent years, arrays of extracellular electrodes have been developed and manufactured to record simultaneously from hundreds of electrodes packed with a high density. These recordings should allow neuroscientists to reconstruct the individual activity of the neurons spiking in the vicinity of these electrodes, with the help of signal processing algorithms. Algorithms need to solve a source separation problem, also known as spike sorting. However, these new devices challenge the classical way to do spike sorting. Here we review different methods that have been developed to sort spikes from these large-scale recordings. We describe the common properties of these algorithms, as well as their main differences. Finally, we outline the issues that remain to be solved by future spike sorting algorithms.

## 1. Introduction

Progress in neuroscience relies to a large extent on the ability to record simultaneously from large populations of cells, in order to understand how information is represented among neurons. One of the most popular techniques to measure such an activity is the use of arrays of extracellular electrodes. With these devices, each electrode records the extracellular field in its vicinity and can detect the action potentials emitted by the neighboring neurons. In contrast to intracellular recording, those extracellular recordings do not give a direct access to the neuronal activity: one needs to process the recorded signals to extract the spikes emitted by the different cells around the electrode. This process is termed *spike sorting*, and many algorithms have been suggested to do it efficiently (see Lewicki (1998) or Rey et al. (2015) for a review).

The first extracellular recordings were performed with a single electrode, and could only give access to 3-5 neurons (Gerstein and Clark, 1964). A recent study (Pedreira et al., 2012) highlighted that the maximal number of accessible neurons should lie between 8 and 10 in that case. Over the last decades, there has been a strong effort to increase the number of electrodes, and therefore the number of recorded neurons. Spike sorting algorithms had to be adapted to process this increasingly large amount of data. At first, electrodes were spaced by hundreds of microns such that the spike of one cell could only be detected on a single electrode (Jones et al., 1992; Shoham et al., 2003). In that case, spike sorting on a large amount of electrodes could simply be done by processing each electrode independently. The parallelization of the problem for large amount of independent electrodes was relatively easy to address.

However, devices where electrodes are packed with a high density have also been developed. The spacing between electrodes is much smaller (tens of microns). As a consequence, a spike from a single cell can be detected on several electrodes. Conversely, each electrode will detect the activity of many cells, a property already encountered in the case of single electrode. This increased density helps a lot to resolve single cells (Gray et al., 1995; Franke et al., 2015a), but electrode signals could not be processed independently. Spike sorting algorithms had to be adapted to this new type of data. While for small numbers of electrodes (e.g. tetrodes), methods that could be seen as adaptations of single electrode sorting worked very well (McNaughton et al., 1983; Harris et al., 2000; Gao et al., 2012), this is not the case with new devices designed with hundreds of electrodes all densely packed. CMOS-based devices with thousands of electrodes have been tested and are now frequently used (Berdondini et al. (2005); Fiscella et al. (2012); Müller et al. (2015); Hilgen et al. (2016)), calling for new algorithmic methods, largely different from the usual sorting methods.

Here we review the different spike sorting algorithms that have been proposed to process recordings from these novel high-density devices. We will first explain the limitations of classical spike sorting approaches to process these large-scale, dense recordings. Then, we will outline the main changes introduced by these new algorithms compared to classical spike sorting approaches. We will emphasize that most of these new methods follow the same global strategy, although they have been developed independently by different groups. Therefore, we will outline the common properties shared by these algorithms, before explaining and discussing their main differences. Finally, we will discuss the issues that still need to be resolved by future spike sorting algorithms.

## 2. The challenge posed by large-scale multielectrode recordings to classical approaches

Most of the classical approaches to spike sorting can be decomposed in two main steps. First, some specific features of the spike waveforms are extracted from the raw data. This allows each spike to be characterized by a small set of numbers/features. Using these features, each spike can now be seen as a point in a low dimension space, and the second step consists in clustering all the points in this reduced space.

For the first step, earliest methods only extracted the spike amplitude (Hubel et al., 1957), and width (Meister et al., 1994) of each spike. More recently, some methods use the full waveform directly when the number of electrodes remains small (Pouzat et al., 2002). Another standard technique is to project each waveform on a set of basis functions (Litke et al., 2004; Quiroga et al., 2004), that are either found by performing a principal component analysis (PCA) on the entire set of waveforms (Egert et al. (2002); Pouzat et al. (2002); Einevoll et al. (2012); Swindale and Spacek (2015)), or by choosing a wavelet basis (Letelier and Weber (2000), Hulata et al. (2002), Quiroga et al. (2004)). For a comparison between PCA and wavelet based analysis, see Pavlov et al. (2007). Note that the two can be combined (Bestel et al., 2012).

Once the dimensionality has been reduced, to tackle the problem of the clustering step, several approaches have been used, but the most standard approach is to fit the clusters with a mixture of Gaussians (Wood et al., 2004; Rossant et al., 2016; Kadir et al., 2014). However, one could also find in the literature approaches such as paramagnetic clustering (Quiroga et al., 2004), mean-shift clustering (Swindale and Spacek, 2014) or even *k*-means clustering (Atiya, 1992; Chah et al., 2011). Another interesting approach is to consider the most consensual clustering across an ensemble of *k*-means solutions (Fournier et al., 2016).

Not all standard methods strictly follow this workflow. For example, linear filtering is an alternative approach which identifies the optimal linear filter to distinguish one signal, of unknown temporal position but of known waveform, from a finite number of other signals of known waveforms, observed on noisy electrodes. This approach was first proposed by Roberts and Hartline (1975), then by Gozani and Miller (1994) and more recently by Franke et al. (2010). This method is similar to template matching approaches that we will describe later. An alternative approach is independent component analysis (ICA) where the first step demix blindly the data and extract the individual source signals from which spikes are identified (Takahashi et al., 2003; Brown et al., 2001; Jackel et al., 2012). Note that variants, such as the convolutional independent component analysis (cICA) of Leibig et al. (2016), has been developed. However, there is no guarantee that the independent components found by those algorithms are indeed isolated neurons.

While all of these methods can be successful when one electrode captures the signals from a only few cells, and when one cell is only recorded by one or a small number of electrodes, it is not trivial to scale them up to process a large number of densely packed electrodes. In recordings performed by large and dense multi-electrode arrays, the spike waveforms live in a high dimensional space, and this makes the clustering challenging. We will review below some suggested improvements to enable clustering on a large number of electrodes.

Finally, a more fundamental problem with clustering-based approach is that the extraction of features from one spike can be distorted by the presence of other spikes nearby. As a consequence, most of the overlapping spikes are not captured by clustering approaches, because they correspond to points in the feature space that are far from the centers of the corresponding clusters. This is a major challenge for clustering techniques (Bar-Gad et al., 2001), that we will explain in more details below. In large scale and dense multi-electrode recordings, overlapping spikes become the rule rather than the exception. Solving this issue is one of the motivation behind new algorithms, based on template matching, that we will review and discuss in a second part.

## 3. Improvements of the clustering

In order to be able to scale up and perform spike sorting for large number of channels with the classical algorithms mentioned above, several refinements of the clustering have been proposed by various groups.

### 3.1. Improved spike detection

Rossant et al. (2016) have proposed a method that pre-processes the data to make clustering easier for multielectrode sorting. As explained above, the spike of a single cell can be detected on multiple electrodes. Conversely, spikes from several cells can be seen on the same electrode. They designed a flood fill method to group together spikes detected on different electrodes that correspond to a single cell. For this they connect together spikes detected synchronously on adjacent electrodes. The exact algorithm to connect adjacent events bears some similarity with standard image processing algorithms, like the Canny contour detection. Spikes are therefore defined as spatio-temporal events, with a given spatial extent, called a mask, for each of them.

In a second step, for each of these events, they remove any voltage deflection outside of the mask, and replace it with noise. This masking removed part of the distortion induced by other spikes when estimating the features, and improved the performance of the clustering. While this improvement is of great help for overlapping spikes that are distant enough in space, it is less clear how it will help for spikes coming from two cells that are physically close. In that case, some electrodes will detect spikes from the two cells, and their masks will strongly overlap. Therefore this masking process may only help avoiding temporally overlapping spikes from distant cells.

### 3.2. Pre-clustering

Marre et al. (2012) and Swindale and Spacek (2014) use a method to break down the clustering problem into multiple smaller parts. After detecting all the spikes in the recording, waveforms are grouped in different subsets according to the electrode where the highest voltage peak was found. Instead of performing a single clustering algorithm on all the waveforms, this grouping outputs *N* subsets, if *N* is the number of electrodes. Each subset contains all the spikes peaking on the same electrode. A clustering is then performed on each of these subsets independently.

Note that this pre-grouping does not assume that the spikes are only detected on a single electrode, which would amount to multiple single electrode sorting. Here, after this pre-grouping, a clustering is performed for each group, and this clustering used the information available on all the electrodes. This simplification allows reducing drastically the number of spikes that have to be processed together. It also allows a simple parallelization of the clustering, which is crucial for large-scale recordings with hundreds or thousands of electrodes.

The main issue with this method is that a cell that is located between two electrodes might emit spikes that peak alternatively on one or the other electrode. In that case, the cell will be split between two different groups, and subsequently in two different clusters. This strategy has therefore to be combined with a later step where all the clusters that correspond to the same cell are merged together. This method is therefore on the side of overclustering the spikes, and merging the different clusters later on. However, merging clusters is usually easier than splitting them since there is one possible result for the first operation whereas the second one presents many possible solutions.

### 3.3. Main issues associated with clustering

A complete review of all the clustering algorithms used for spike sorting is beyond the scope of this review. However, we would like to outline the main issues associated with the clustering step, that are common to almost every clustering algorithm.

#### 3.3.1. Mathematical definition and non-linear optimization

Two of the main issues associated with any spike sorting solution relying on a clustering approach can be found in the roots of the clustering *per se*. Mathematically, the clustering suffers from a lack of problem statement and problem resolution. First, one need to agree on a mathematical definition of the notion of cluster to state the problem. Because there exists many different cluster models (e.g. centroid models, distribution models, density models), there are numerous notions of what a cluster is. It is not obvious if one of these notions would fit appropriately to the biological reality. Hence, the first problem is that the stated problem is an approximation of the true problem. Thus, the solution to this clustering problem is an approximated solution to the true problem. This is why it often requires the user to spend a rather large amount of time in manual curation, because the solution to the true problem is in the neighborhood of the approximated solution.

Second, solving a clustering problem brings additional issues. The different methods used to do clustering involve finding the minimum of an objective function, and the solution landscape almost surely presents local minima. As a consequence, running twice the same clustering algorithm with two different set of parameters (i.e. internal parameters such as initial centroids for the *k*-means algorithm) can lead to different results. The reason is that the two runs can be trapped in two different local minima. In many cases it takes several trials before converging to the global minimum, which increases the computational cost. In practice, the algorithm may stop before convergence because the more complex/challenging is the solution landscape, the less likely is the convergence in a reasonable time.

#### 3.3.2. Overlapping spikes

More importantly, as mentioned above, a major issue with clustering is that it will miss many overlapping spikes. If two spikes are overlapping on the same electrode, there will be a distortion in the feature estimation, that will drive the spike beyond the limits of the cluster defined on isolated spikes. Note that the superposition problem has been known for a long time (Prochazka et al., 1972; Roberts and Hartline, 1975). The issue was apparent in Harris et al. (2000): they showed that the error rate of the spike sorting is strongly increased during spindle waves, which are epochs of synchronous firing. False positive errors could change from 5% to almost 80%, and false negative were also increased by at least 20%. The issue was more extensively studied by Pillow et al. (2013), where they show that synchronous spikes will be missed by a pure clustering approach. An additional study of Franke et al. (2015a) confirms that clustering-based methods perform poorly for overlapping spikes, as shown by Lewicki (1998) and Quiroga et al. (2004). Template matching approaches have been developed in order to deal with these overlapping spikes.

## 4. Template matching approaches

Several template matching approaches have been developed for spike sorting (Pillow et al., 2013; Pachitariu et al., 2016; Marre et al., 2012; Yger et al., 2016; Prentice et al., 2011). Note that historically the use of template matching (Gerstein and Clark, 1964) predates the use of clustering (Simon, 1965) and then experienced renewed interest. All these methods usually assume that the extracellular signal can be decomposed as a sum of so-called “templates” (one template is the average extracellular waveform triggered by one neuron) plus some noise:

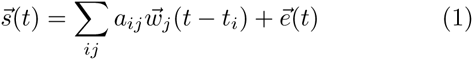

where 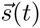 is the signal recorded over the electrodes of the multi electrode array and over multiple time points. 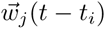 is the spatiotemporal template associated with each cell, which represents the average waveform triggered on the electrodes by cell *j* (example in figure 1B). *t*_*i*_ are all the putative spike times over all the electrodes, *a*_*ij*_ is the amplitude factor for spike time *t*_*i*_ for cluster *j*, and 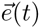 is the background noise.

In this notation, the spike train associated with cell *j* is the set of times *t*_*i*_ where *a*_*ij*_ is different from zero. The template matching approach aims at finding the right values for 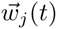 and *a*_*ij*_, i.e. to find where each cell spiked. Almost all the template-matching based methods try first to find the value of the templates, and then the values of *a*_*ij*_. Depending on the algorithm, the amplitude values can only be 0 or 1, or can take any continuous value. We will review these methods in subsections 4.3 and 4.4.

### 4.1. Template extraction

To estimate the templates, most methods usually rely on clusters extracted from the recording using one of the methods described above. Each cluster corresponds to a set of snippets in the extracellular data. The snippets of a given cluster are supposed to be realizations of action potential of a single cell. We want to extract a canonical representative (i.e. a template). A naive method would be to consider the average waveform of these snippets. However, because averaging is very sensitive to outliers, if some of the snippets also include overlapping spikes from other cells, they might distort the estimate of the template. Two solutions have been developed to circumvent this issue. The simplest (and fastest) one is to take the median at each time point instead of the mean (Marre et al., 2012; Yger et al., 2016). The median is way less sensitive to outliers than the mean. This method usually solves the issue of overlapping spikes.

Another solution is to model the extracellular signal from the clustering result:

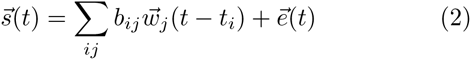

Notations are similar to equation 1, except that *b_ij_* are binary variables such that *b_ij_* is set to 1 if *t_i_* is associated to cluster *j*, and to 0 otherwise. Here the unknown variables are the templates 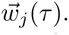 Under these conditions, it is possible to find the templates that will fit the extracellular data best, by minimizing the following square difference (Pillow et al., 2013; Ekanadham et al., 2014):

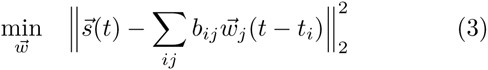

The two methods seem to give similar results^1^ although, in theory, the first approach is less sensitive to noise, whereas, the second one is less sensitive to strong correlations between cells (i.e. overlapping spikes). This is due to the fact that taking the median is a way to minimize the *ℓ*1-norm between the different snippets and the template, while equation 3 is a minimization of a *ℓ*2-norm.

### 4.2. Finding the spike trains

Once the templates are found, we need to find when they appear on the extracellular signal. For this, template matching methods usually use algorithms similar to projection pursuit (Friedman and Tukey (1974), although with different criteria for acceptance and stop). Most of them can be summarized as an iterative greedy approach with the following steps, for a given time chunk (illustrated in figure 1A):

**Figure 1:**
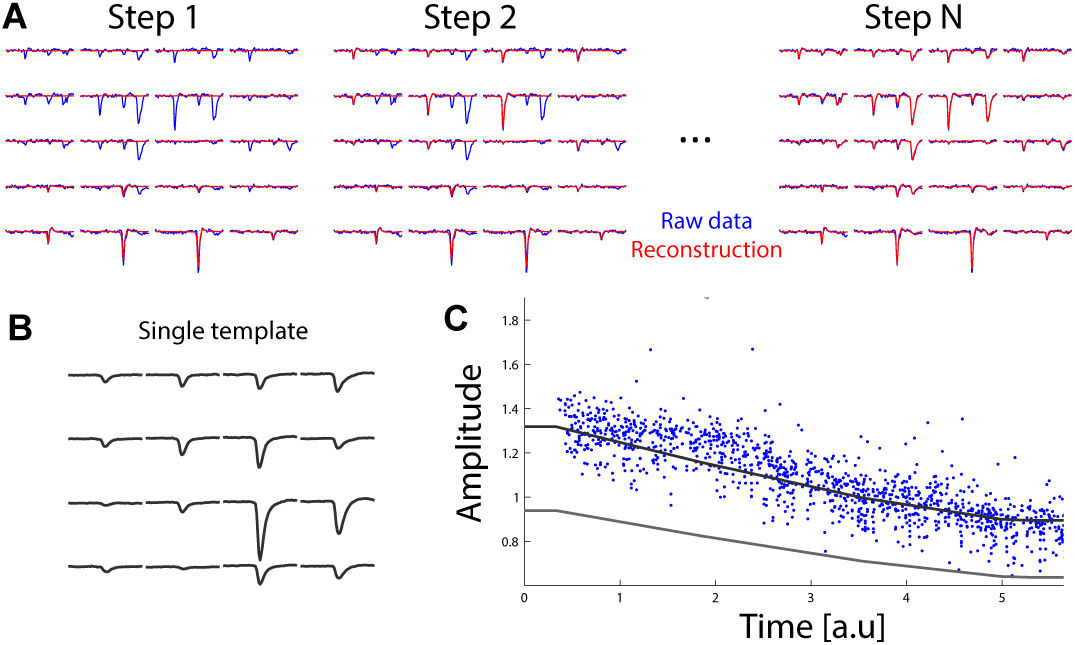
The template matching approach. **A:** illustration of the iterative template matching approach. The extracellular signal (in blue, shown for 20 electrodes) is matched iteratively with a sum of templates. At each step, a template is added to the signal (red) to match better the data. At the end, all the spikes are fitted by a template, and the sum of templates (red) predict very well the data (blue). **B:** example of a single template over 16 electrodes. **C:** example of amplitude values fitted to the data for one template, as a function of time. Gray lines represent the average amplitude over time, and the minimal amplitude over time (see text for details).

1. Find the template that matches best the raw data. If amplitude is allowed to be different from 1, find the best matching amplitude.
2. Define a criterion to accept the template. It can either be about the quality of the fit to the raw data, or about the value of the best amplitude, or both.
3. If the template is accepted, subtract it from the raw data. Then go back to the first step.

The different algorithms that have been proposed differ mostly in the acceptance criterion, and in the possibility to have amplitude different from 0 and 1 or not.

One common issue that needs to be mentioned before comparing the approaches is sampling jitter. When a cell emits a spike, the spike time may peak at a time *t* + *dt*, where *t* is the closest time point sampled by the data acquisition, and *dt* is the time difference between the true spike time and *t*, smaller than the acquisition period. As a result, in template matching approaches, a template will be matched at time *t* to explain a spike that occurred at *t* + *dt*. The compensation of this *dt* is necessary (McGill and Dorfman, 1984) when one does not use a high sampling frequency. For example, Prentice et al. (2011) use linear interpolations, Pillow et al. (2013) use local approximations based on Taylor expansions and Yger et al. (2016) use similar expansions (see also Marre et al. (2012) where this issue is mentioned). Additional solutions, such as polar expansions, were developed by Ekanadham et al. (2011).

### 4.3. Approaches with binary amplitudes

Segev et al. (2004), Pillow et al. (2013) and Franke et al. (2015b) assume that the amplitude of a template is always equal to 1 (*a_ij_* ∈ {0,1} in equation 1). Segev et al. (2004) keep a template if it improved the prediction of the extracellular signal by the sum of templates, i.e. if subtracting it to the raw data led to a reduction in variability that passes a given threshold. This threshold is needed to avoid overfitting the noise with small templates. Pillow et al. (2013) base the criterion of acceptance on an objective function: the value of the function had to be improved when fitting an additional spike. This function is the sum of two terms:

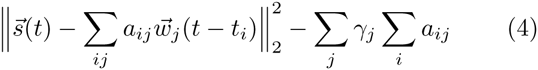

The first one is the square difference between the extracellular signal and the sum of templates, in the metric defined by the noise covariance. It will usually decrease if an additional spike is fitted to the signal. The second term is a regularization on the average firing rate of each cell, and corresponds to a cost per spike. This term decreases when an additional spike is fitted to the signal, and reflects the prior that cells are more likely to be silent (i.e. respect their firing rate) than to fire all the time. This second term is here to avoid overfitting the noise with small templates. Note that, while this term is called a prior by Pillow et al. (2013), it is based on the data (on the measured firing rate for each cell). We will call it a regularization term in the following. Conceptually, we can see that the two methods are quite similar. If we whiten the extracellular signal before template matching, then the first term in the objective function of Pillow et al. (2013) is equivalent to the square difference between the extracellular signal and the sum of templates, which is exactly what Segev et al. (2004) use. When Segev et al. (2004) then compare the reduction of this square difference to a threshold, this threshold can be compared to the change of the second term in the objective function of Pillow et al. (2013), which reflects the regularization on the firing rate. The method of Pillow et al. (2013) is more elaborate because the regularization term can change from one cell to the other, while the method of Segev et al. (2004) uses the same threshold for all cells. However, it seems that the exact regularization values does not change much the results of the spike sorting (Pillow et al., 2013). Therefore, we expect that these algorithms should give similar results. More recently, Franke et al. (2015b) used a relatively similar approach but allowed fitting two templates at the same time. This additional feature leads to a better estimation in the case of overlapping spikes.

### 4.4. Approaches with graded amplitudes

Other methods have assumed that a template can be scaled up or down every time the cell spikes: they assume that the amplitude *aij* can take other values than 0 or 1 in equation 1. Prentice et al. (2011) assume that the spike amplitude for a given cell follows a Gaussian probability distribution, whose mean is equal to 1. The standard deviation of the distribution is estimated from the previously found cluster. Then, they maximized an objective function that has two terms: the first one is the same as the one of Pillow et al. (2013), i.e. the difference between extracellular signal and the sum of templates in the noise covariance metric. The second one is a regularization term that reflects two facts. First, an amplitude closer to 1 is more likely than a very small, or a very big one. Second, a template with a high firing rate is more likely than another one with a low firing rate. By balancing these two terms, the optimization process avoid to add a lot of templates with small amplitudes that are highly unlikely. It also avoid to add a lot of templates associated to units with low firing rates. This second term can thus be understood as a combination of two regularization constraints: one over the amplitudes, and another one over the firing rates.

Marre et al. (2012) and Yger et al. (2016) also allow amplitude variations, but the acceptance criterion was different: after having found the amplitude that best matches the extracellular signal, the template was kept if the amplitude was between thresholds, *a_min_* and *a_max_*. At first sight, this criterion seems surprising since it does not depend on the improvement in the quality of the fit. In fact, the process of finding the best amplitude is by itself an estimation of the improvement of the fit. For a given iteration, if we note 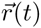 the extracellular signal that remains to be fitted (i.e. after subtraction of the templates fitted in the previous iterations), and 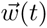 the new candidate template that needs to be fitted, then the best matching amplitude *a* will be found by minimizing 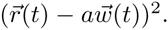 It can be shown that, if this template is accepted, the square difference will decrease by 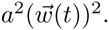 In other algorithms, this decrease of the square difference has to be larger than the increase of the regularization term for the template to be accepted. Setting a minimal amplitude is therefore equivalent to having a regularization term that is different for each cell, but constant as a function of the amplitude of the spike (similar to what was done by Pillow et al. (2013)). The other threshold for maximal amplitude is less important, and only plays a role to avoid very high, unrealistic values.

The advantage of having an amplitude threshold as a parameter, instead of a threshold for improvement in the goodness of fit, is that this parameter is much more intuitive for the user: we can figure out reasonably well what a minimal amplitude of 0.4 or 0.8 means. Thresholds on goodness of fit are less easy to understand. Furthermore, by looking at the set of amplitudes fitted over time, we can get a sense of the right values for these amplitude thresholds. If the minimal amplitude threshold is too low, the template is also fitted on noise, with small amplitudes which are clearly different from the amplitude of real spikes, that are close to 1. When we labeled the pairs of spikes with refractory period violations, we often see that most of them involve one of these spurious fits. It is therefore easy to readjust the threshold to a correct value. These thresholds can also be made time dependent, as can be seen in figure 1C. This gives more flexibility to process non-stationary data while keeping understandable parameters. Of course, the disadvantage of this method is that the algorithm is not expressed as the minimization of a cost function.

### 4.5. Different algorithms correspond to different assumptions about spike amplitude distributions

Can all these methods be expressed with an objective function having a similar structure? As we showed before, all the three methods discussed above (Prentice et al., 2011; Marre et al., 2012; Pillow et al., 2013) aim at minimizing the square difference between the extracellular signal and the sum of fitted templates. The difference lies in the regularization term, which reflects an assumption about the possible amplitude for the spike. More formally, the quantity we want to minimize is:

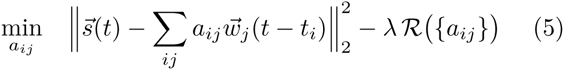

where the first term is the square difference between the data and the reconstruction model, *R* denotes the regularization function over the amplitude values *a*_*ij*_ and *λ* is a free parameter (i.e. trade-off between the two terms).

In Prentice et al. (2011), the regularization term reflects an assumed Gaussian distribution for the amplitude. In Marre et al. (2012), the amplitude thresholds might reflect an assumption of flat amplitude distribution between the minimal and maximal amplitudes, and 0 elsewhere.

With a similar approach, Ekanadham et al. (2014) use a *ℓ*1-minimization algorithm to find the right amplitudes. This *ℓ*1-minimization is equivalent to assuming that the spike amplitude distribution that has the form of a power law, i.e. 1*/*(*ϵ* + *a*)^*p*^. The form of this distribution gives an advantage to small amplitudes. As a consequence, this algorithm outputs a lot of small amplitude spikes, and this is later corrected by removing all the spikes whose amplitude is smaller than a given threshold. The threshold is estimated *a posteriori* by fitting a Gaussian distribution to the amplitude distribution found empirically.

One way to summarize the difference between these three methods is therefore to say that they differ in their assumption on the amplitude distribution. Prentice et al. (2011) assume a Gaussian distribution, Marre et al. (2012) and Yger et al. (2016) assume a flat distribution between some thresholds, Ekanadham et al. (2014) and Pachitariu et al. (2016) assumed a power-law distribution in the core of the algorithm, but corrected it later on with a Gaussian distribution.

### 4.6. Caveats when minimizing an objective function

While this is an intuitive way to explain the differences between the different algorithms, it has to be noted that some sorting algorithms do not directly minimize the objective function described above. For example, in both Prentice et al. (2011) and Marre et al. (2012), during the iterative process, the amplitude was chosen as the one that best matches the data, without taking into account the regularization term on the amplitude values. Formally, the amplitudes were chosen to be the solution of:

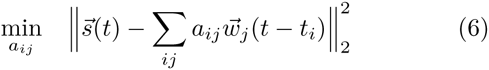

A direct minimization of the total objective function, including the regularization term, would have biased all the amplitudes towards 1 since the amplitudes would have been the solution of:

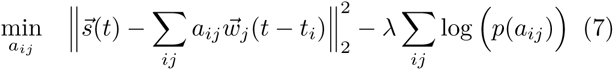

where *λ* is a free parameter and *p*(*a*_*ij*_) is the probability density function of the amplitude values. This bias affects the quality of the fit, and can lead to fitting other templates where templates have been fitted with biased amplitudes. For example, figure 2A shows a comparison of several errors function used while optimizing the amplitude of a given waveform, displayed in figure 2B. As we can see, the choice of the error criteria can have a strong impact on the “optimal” amplitude, leading to more or less pronounced residuals (see figure 2C). To avoid this, it is necessary to take the amplitude value that best matches the data, without any regularization, and only use the regularization to decide afterwards whether this template should be kept or not.

**Figure 2:**
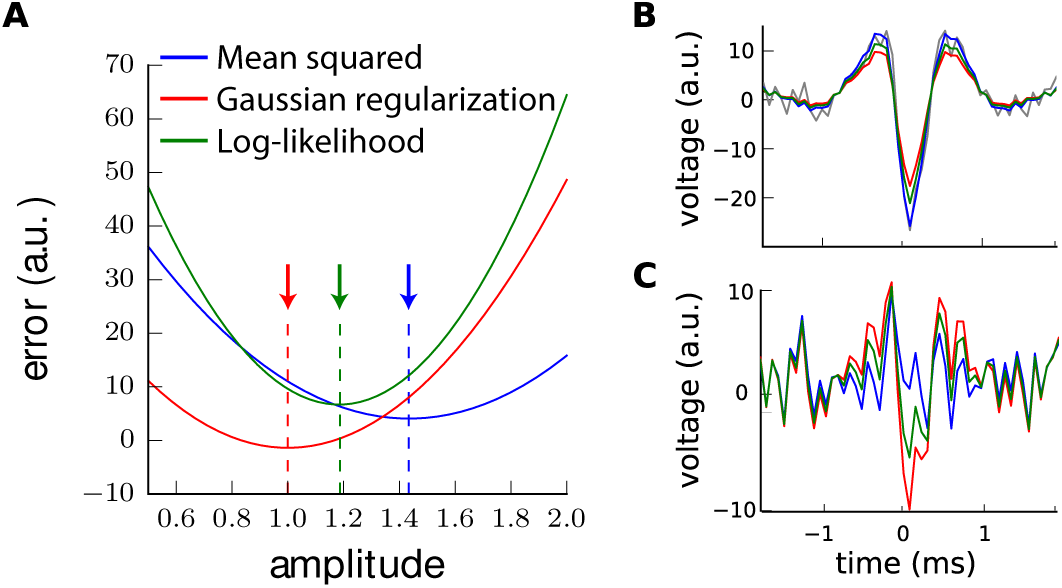
Illustration of biased amplitudes toward 1 when minimizing the log-likelihood. **A**: Comparison of the error function used for the optimization of the amplitudes. Mean squared error of the residual, as described in equation 6 (blue). Penalty which comes from a regularization with a Gaussian distribution on the amplitude values (red). Log-likelihood, as described in equation 7 (green). The dotted vertical lines indicate the minimium of each of these error functions. **B**: Illustration of the results of the fit, with optimal scaled waveforms for each error function superimposed onto the raw data (gray), colorcoded as in **A**. **C**: Residuals (fit minus raw data) for each of those error functions, colorcoded as in **A**.

### 4.7. Assumptions behind the template decomposition

An important question is whether template matching algorithms can always replace clustering algorithms, or if they have some intrinsic limitations that make them less flexible than clustering. This is still an open question, and only direct comparisons between the different approaches, in cases where the true solution is known, will tell us what is the best approach (Yger et al., 2016). Here we would like to give some intuition about how the main assumptions of template matching approaches can translate in the feature space that clustering approaches use.

In the template matching approach, the noise is supposed to be independent of the templates. In the case where no amplitude variation is allowed, it means that the variability in the snippets always comes from the same noise source. In a given feature space, it means that all the clusters should be elongated in the same directions. This is illustrated in figure 3A: while the clusters have different centers, they are all ellipses extended in the same directions.

If the spike amplitude is allowed to change, this means that, in a feature space, each cluster has two sources of variability: a common one, which corresponds to the noise, and another one that is specific to each template. The second one is constrained to be in the direction of the template, which is approximately the cluster center. Therefore, in a feature space, it means that the clusters have now noise in common directions, but also an elongation that will follow the arrow that connects the point 0 in the feature space, and the center of the cluster (figure 3B). This is more realistic than the previous assumption, but it is not clear whether this gives a good account of all the variability found for each cluster.

**Figure 3:**
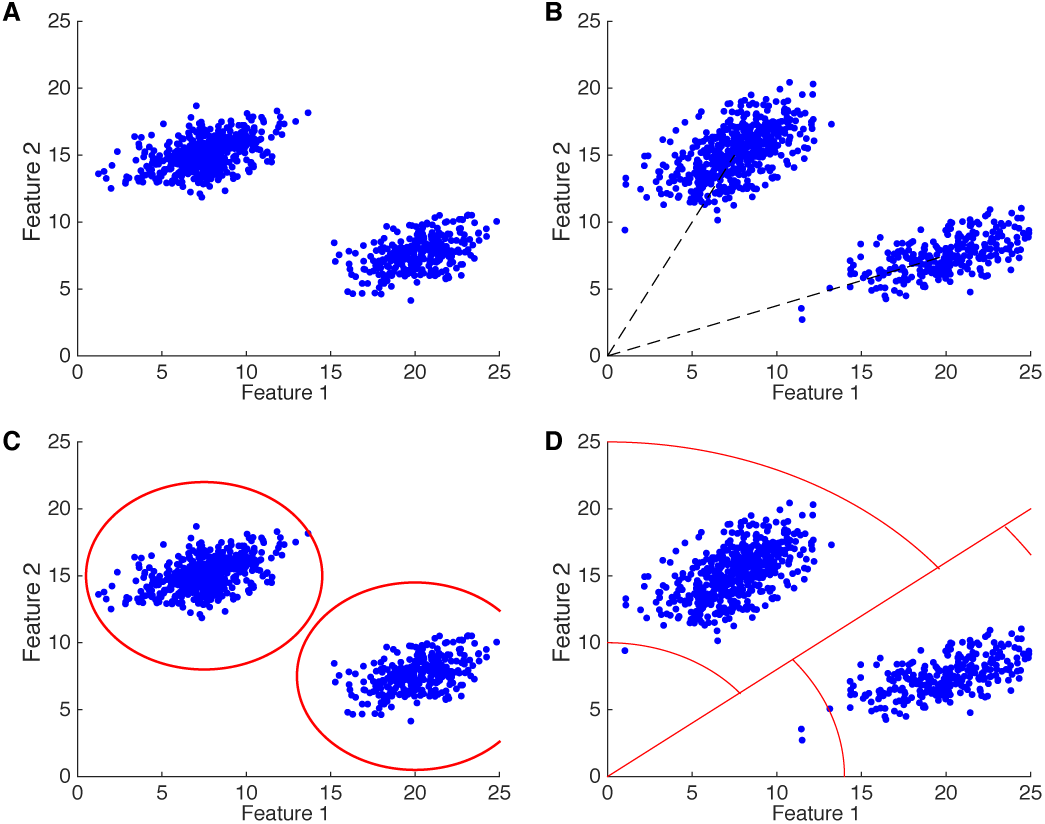
Illustrations of the assumptions of the template matching in the clustering space. **A**: Example of two clusters in the feature space, when assuming that they are generated by templates with no amplitude variation. **B**: Same than A, but now with the assumption that the template can vary in amplitude according a Gaussian distribution. **C**: Equivalent borders (see text) for the clusters for a template matching that chooses the template closest to the spike. **D**: Equivalent borders in the case where the template is chosen based on the spike shape, and that only a certain range of amplitude is allowed. See text for details.

If we were to use template matching only on isolated spikes, we could also define areas in the feature space where a point is assigned to a given template. A snippet is always assigned to the best matching template. In some algorithms (Pillow et al., 2013), it means this template is closest in the sense of the least square difference. In the feature space, this means that a point will always be assigned to the closest centroid. We can use this rule to define equivalent cluster borders (figure 3C). In other algorithms (Prentice et al., 2011; Marre et al., 2012), only the spike shape is used to define the best matching template, and then the algorithm decides whether the best matching amplitude is plausible or not. This defines different shapes for the border: a straight line from the 0 point to separate the regions of preferred spike shape and some circles to define the allowed amplitudes, following the approach of Marre et al. (2012). Figure 3D illustrates these shapes.

So the competition between the different templates defines some natural borders. There is no guarantee that this is the best and proper definition for the cluster borders. Future works will need to address this issue by comparing the results of template matching and clustering algorithms. However, the intuitions we have drawn here can be used to compare more intuitively pure clustering-based versus template matching approaches.

## 5. Conclusion: challenges ahead

The methods described here have enabled to sort spikes from a large number of cells and electrodes (Yger et al., 2016; Pachitariu et al., 2016). However, there are still several challenges that need to be overcome. First, most of the algorithms described here have been tested on *in vitro* data, in the retina (but see Ekanadham et al. (2014), Frankeetal. (2015b) or Ygeretal. (2016) for *in vivo* tests). *In vivo* tests on silicon probes with a large number of recording sites close apart will be necessary. A possible required improvement is a better description of the cluster (Yger et al., 2016). As we explained above, the template matching makes some assumptions about the shape of the clusters, and it is not clear if these assumptions are verified or not *in vivo*. A related issue with spike sorting is the need to have more ground truth data, i.e. recordings where at least one cell is recorded with another technique, so that we know when the spikes occur. These data are essential to test spike sorting algorithms (Neto et al. (2016)).

A second point is that template matching does not replace clustering. All the methods described require a set of clusters, from which the templates can be extracted. The clustering can do mistakes that can be tolerated, as long as they do not distort the template estimation. But a decent performance in clustering is nonetheless required. So one still needs an efficient way to cluster. Ekanadham et al. (2014) and Pillow et al. (2013) have proposed to do back and forth between template estimation and finding the amplitudes. This is an extension of the approach we described previously: after finding the amplitudes, they are used to estimate the templates again with a least square method. Then this new set of templates is fitted once again to the data. Note that this global iteration does not remove the need for an initial clustering, so that the templates are properly initiated (at the very least, they need to be in sufficient numbers). The interest of doing multiple iterations of template estimation and matching is not completely clear. While Ekanadham et al. (2014) claim that it is crucial, Pillow et al. (2013) mention that there is only a marginal improvement after the first pass. Another modification of the iterative approach can be found in a work of Franke et al. (2015b), where solutions beyond this iterative approach have been developed that can lead to a better sorting of synchronous spikes.

Another challenge is the time spent on manual curation. Even the best clustering makes mistakes, and some cells will be represented by more than one template. Finding all the pairs that need to be merged require a significant amount of time for hundreds of electrodes. Methods need to be developed to make this kind of tasks as automated as possible, so that the time spent by the user is reduced to a minimum (Yger et al., 2016).

Nowadays, new devices with CMOS components now allow recordings from thousands electrodes simultaneously (Berdondini et al. (2005); Fiscella et al. (2012); Müller et al. (2015); Hilgen et al. (2016)), and it remains to be seen it these algorithms can scale up and process such a large amount of data. We need to be sure that the time spent on manual curation can remain small enough that we can get thousands of spike trains in a decent amount of time (see preliminary evidence that it might be the case by Yger et al. (2016)).

Finally, one problem that needs to be properly tackled by the new generation of spike sorting algorithms appears during long lasting chronic recordings (Nicolelis et al., 2003). It is indeed well known that because of tissue changes, or because of experimental protocols, recordings can be non-stationary and drifts in the neuronal waveforms can appear over long time scales. For any template matching based approach, one should rather consider spatio-temporal kernels that could evolve over time, and be distorted. To some extent, some of these deformations can be dealt with by allowing graded amplitudes for the templates (see for example figure 1C, where the amplitude evolves over time). However, a more robust framework is required for a better understanding of the drifts, especially because latest algorithms (Yger et al., 2016; Pachitariu et al., 2016) seem to pave the way toward real-time spike sorting. Such an understanding would be crucial in the context of accurate online spike sorting.

## 6. Acknowledgments

This work was supported by ANR OPTIMA and TRAJECTORY, the French State program Investissements d’Avenir managed by the Agence Nationale de la Recherche [LIFESENSES: ANR-10-LABX-65], a grant from the European Union Seventh Framework Programme (FP7/2007-2013, grant agreement no. 604102, Human Brain Project), and NIH grant U01NS09050 to OM.

## Footnotes

1 See http://phy.cortexlab.net/data/sortingComparison/ for a direct comparison on some synthetic ground-truth datasets

